# Deep learning-based interpretable prediction of recurrence of diffuse large B-cell lymphoma

**DOI:** 10.1101/2024.06.03.596955

**Authors:** Hussein Naji, Juan I. Pisula, Stefano Ugliano, Adrian Simon, Reinhard Büttner, Katarzyna Bożek

## Abstract

**Background:** The heterogeneous and aggressive nature of diffuse large B-cell lymphoma (DLBCL) presents significant treatment challenges as up to 50% of patients experience recurrence of disease after chemotherapy. Upfront detection of recurring patients could offer alternative treatments. Deep learning has shown potential in predicting recurrence of various cancer types but suffers from lack of interpretability. Particularly in prediction of recurrence, an understanding of the model’s decision could eventually result in novel treatments.

**Methods:** We developed a deep learning-based pipeline to predict recurrence of DLBCL based on histological images of a publicly available cohort. We utilized attention-based classification to highlight areas within the images that were of high relevance for the model’s classification. Subsequently, we segmented the nuclei within these areas, calculated morphological features, and statistically analyzed them to find differences between recurred and non-recurred patients.

**Results:** We achieved an f1 score of 0.83 indicating that our model can distinguish non-recurred from recurred patients. Additionally, we found that features that are the most predictive of recurrence include large and irregularly shaped tumor cell nuclei.

**Discussion:** Our work underlines the value of histological images in predicting treatment outcomes and enhances our understanding of complex biological processes in aggressive, heterogeneous cancers like DLBCL.

## 1 Introduction

Diffuse large B-cell lymphoma (DLBCL) represents the most prevalent form of non-Hodgkin’s lymphoma (NHL) in adults, accounting for 30%-40% of all NHL cases. It originates from B-lymphocytes and is characterized by its remarkable diversity in terms of morphology, biology, and clinical outcomes. Despite treatment strategies that include high-dose chemotherapy coupled with autologous stem cell transplantation, achieving long-term remission remains challenging. Approximately half of all patients experience a recurrence, following initial therapy. Due to its aggressiveness, recurred patients have a critically low survival probability [1]. Thus, there is a pressing need for a reliable upfront detection of those patients that will not benefit from standard frontline regimens.

In the recent past, deep learning has made remarkable progress in predicting recurrence of various cancer types (including bladder [2], breast [3-6], colorectal [7,8], hepatocellular carcinoma [9,10], lung [11,12], and prostate cancer [13,14]) based on histological images.

Previous studies have used deep learning models (mostly variations of ResNet architectures [15]) to predict recurrence based on images as well as clinical features such as tumor grade, stage, age, etc. [2-5,9,11]. While offering good performance, these models did not provide insights into the determinants of the patient outcomes.

Explainability of recurrence prediction models is of tremendous medical importance. Identifying which image regions are predictive of specific clinical outcomes could potentially lead to discovery of novel visual biomarkers as well as to a better understanding of the underlying resistance mechanisms. Several recent studies have included interpretability analyses using multiple instance learning (MIL) [12,13] or gradCAM heatmaps [6] to highlight image patches that were important for the model’s decision. Although these methods allowed for qualitative insights into the relevant image regions, they did not perform an in-depth quantitative analysis of those regions.

There is relatively little research on the image-based prediction of DLBCL patient outcomes. Up to 2019, the primary prognostic features were clinical features, gene expression markers, and genetic abnormalities [16]. Wang et al. [17], Fan et al. [18], and Xing et al. [19] used traditional machine learning algorithms (such as random forest support vector machine, regression, gaussian mixture model clustering) to predict recurrence in patients with DLBCL using clinical features. Shankar et al. [20] introduced LymphoML, an interpretable deep learning method that identifies morphological features that are predictive of lymphoma subtypes. Their model points to concrete single-cell-based morphological features that are distinctive for different lymphoma subtypes, such as minor axis length and nuclei area. However, how these features could be leveraged to predict recurrence in DLBCL patients is an open question.

In this work, we address the challenge of prediction of recurrence after first-line treatment in DLBCL and interpretation of the prediction model. We developed a deep learning-based prediction pipeline based on tissue microarrays (TMAs) stained with hematoxylin and eosin (H&E). We extracted the model’s high attention areas of the core images, applied nuclei segmentation on these areas, and determined single-cell morphological features which are distinct for recurred and non-recurred patients. While the causes of DLBCL relapse are still unknown, here, we point to cell morphology features which might be determinant for patient response to treatment.

## 2 Methods

### 2.1 Patient Cohort and Dataset

We use a publicly available dataset of the Stanford Cancer Institute [21]. The data is based on a study cohort with 167 patients, containing 306 cores (1-2 cores per patient) from seven H&E-stained TMAs. The TMA slides were scanned at 40x magnification (0.25 μm per pixel) on an Aperio AT2 scanner (Leica Biosystems, Nussloch, Germany). Each TMA has a thickness of 4 μm and includes a formalin-fixed, paraffin-embedded (FFPE) section of tumors arranged in a grid. Each core within the TMAs has a diameter of 0.6 mm [21]. The clinical data shows that 47 patients of the cohort did suffer from a recurrence, whereas 120 patients stayed recurrence-free.

### 2.2 Pipeline

We developed a pipeline (Figure 1) that starts with a preprocessing step (Figure 1A) in which the cores are cut into non-overlapping patches of size 256 x 256 pixels. For each patch of a core, a low-dimensional feature representation is determined using a ResNet50 model pretrained on ImageNet. This results in a 1024-dimensional feature vector which is then further compressed to a 512-dimensional vector (Figure 1B) through linear transformation of a fully connected layer. To predict recurrence from the images, we use CLAM (clustering-constrained attention multiple instance learning) [22]. CLAM leverages attention-based learning to localize image regions that are of significant diagnostic importance. That is, the model uses an attention-based pooling function to aggregate patch-level features into sample-level representations to classify a sample into ‘recurrence’ or ‘non-recurrence’. The classification decision depends on the attention score that is assigned to each patch and indicates its importance to the sample-level representation for the respective class (Figure 1C) [22]. For training the classification model, we used 60% of patients for training, 20% for validation and 20% for final testing. We further applied a cross validation with 4 iterations to further validate our model’s performance. Hence, we express the performance as the mean of all iterations. Out of the correctly classified cores (true positives), we utilized the top 20 patches with the highest attention scores for further analysis (for both groups, respectively). We segmented the nuclei within each patch using a modified version of HoVer-Net [23], that has specifically been trained on H&E-stained DLBCL whole slide images (Figure 1D) [24]. We excluded cut-off nuclei at the patch margins, quantified morphological features of the remaining segmented nuclei, and applied statistical analysis to identify differences between the features of recurred and non-recurred patients (Figure 1E). We quantified the following shape- and size-based nuclei features:

**Figure 1.**
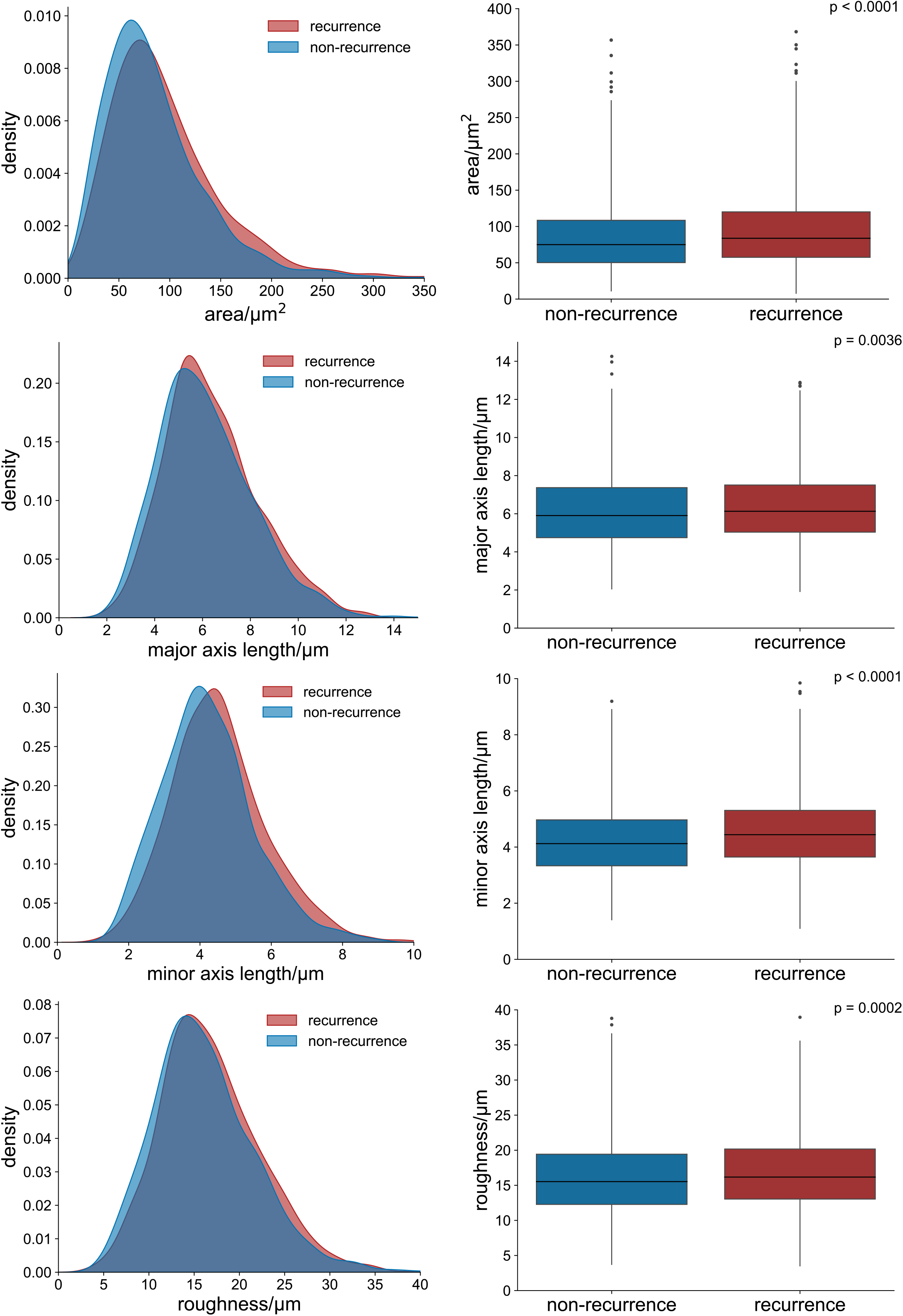
Deep learning pipeline. **A** H&E-stained core images were segmented and cut into non-overlapping patches of size 256 x 256 pixels. **B** Features were extracted from all patches of a core image using a ResNet50 encoder. The final feature vectors had a size of 1024. **C** CLAM model to classify the cores into ‘recurrence’ or ‘non-recurrence’. The model assignes attention scores to each patch of a core image, with blue frames indicating low attention patches and red frames indicating high attention patches. **D** Segmentation of the nuclei of the top 20 patches (based on the attention score) of each class using a pre-trained modified HoVer-Net architecture^24^. **E** We calculate morphological features from the segmented cell nuclei and statistically analyze them to find significant differences between recurred and non-recurred patients.

- area: area of the nucleus.
- major axis length: length of the major axis of the ellipse that has the same second central moment as the nucleus.
- minor axis length: length of the minor axis of the ellipse that has the same second central moment as the nucleus.
- roughness: variance in the length of a vector that is centered at the centroid of a nucleus as it rotates along with each boundary point. The higher the value, the higher the degree of nucleus shape irregularity [25].

## 3 Results

### 3.1 Relapse Prediction

Our model achieves a classification accuracy of 0.73, a precision of 0.74, a recall of 0.96, and an f1 score of 0.83. A confusion matrix is shown in Figure 2. Despite the class imbalance of the dataset with approximately 28% of patients that experience recurrence, the errors are equally distributed among the two patient groups.

**Figure 2.**
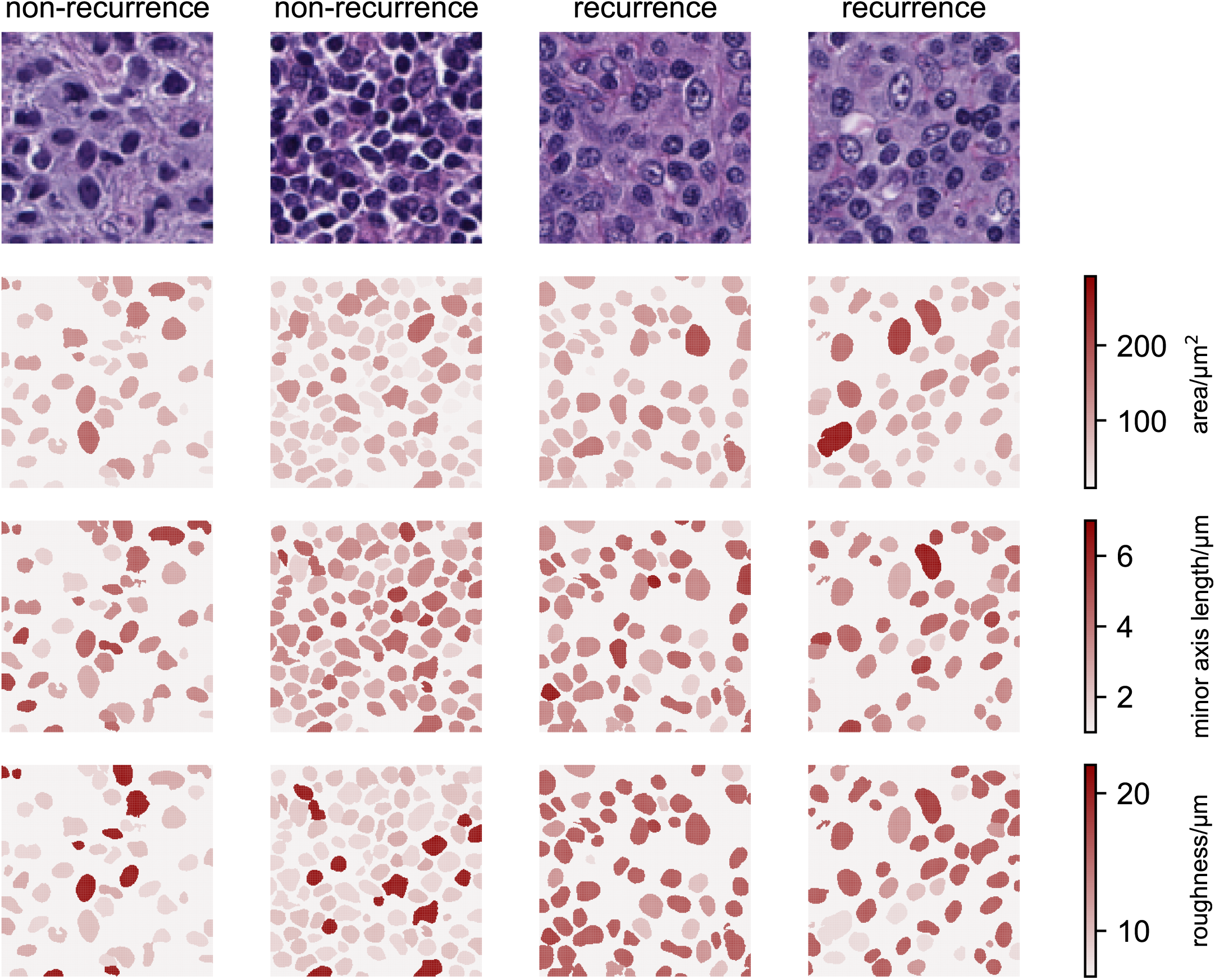
Confusion matrix of the recurrence prediction on the hold-out test set (n=32). The matrix was normalized and shows the average performance across all cross validation folds.

### 3.2 Statistical Analysis

We identified nuclei area, major and minor axis length, and roughness as nuclei features that differ between the patients that have recurred and those that have not (Figure 3). In the high-attention image areas for recurred patients, the nuclei area, major and minor axis length, and roughness are higher, with a difference of 10.8 μm^2^ (p < 0.0001, t-test), 230 nm (p = 0.0036, t-test), 321 nm (p < 0.0001, t-test), and 706 nm (p = 0.0002, t-test), respectively.

**Figure 3.**
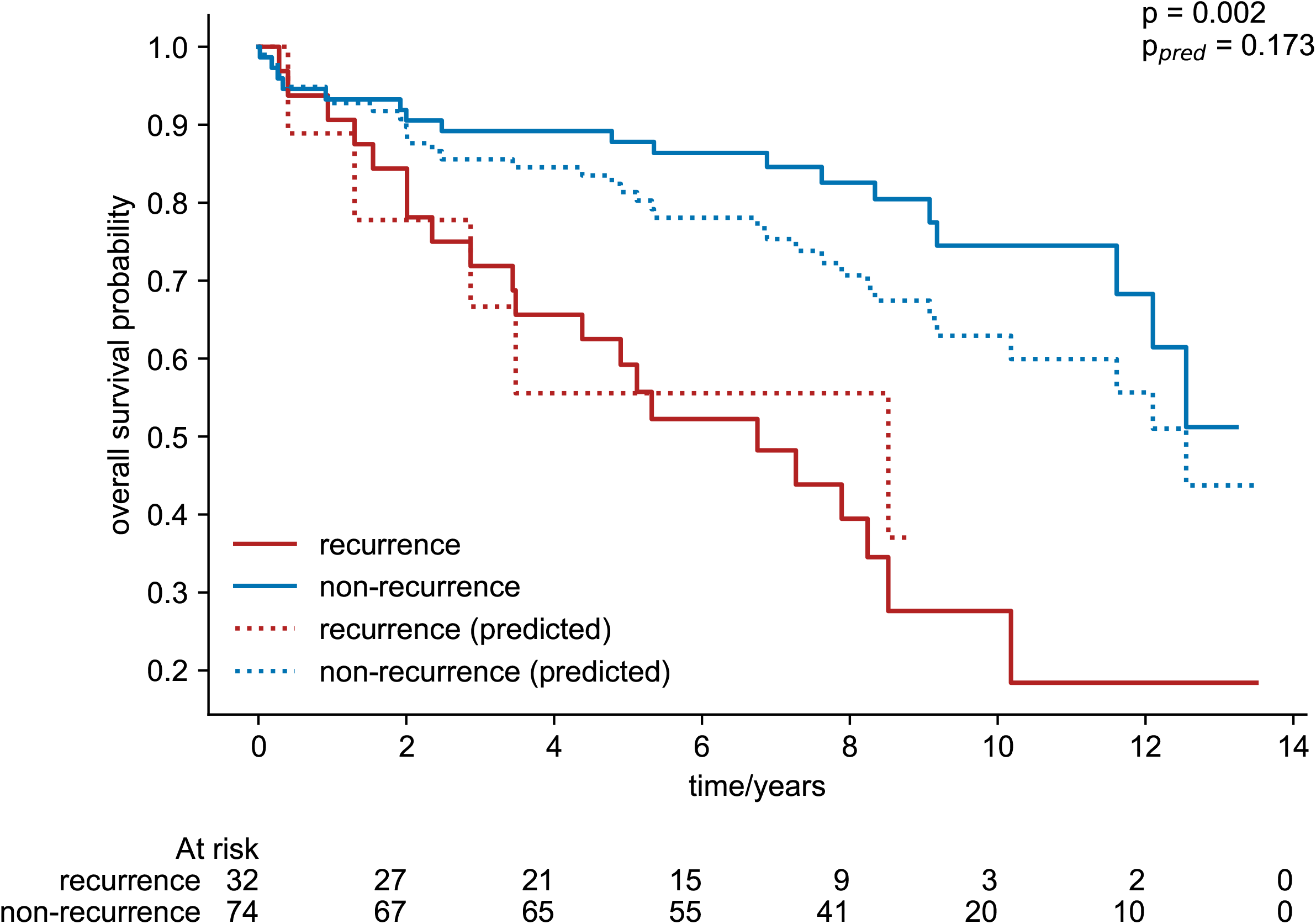
Distribution and boxplots of the determined morphological features for recurred (n=18, with an average of 63 nuclei per patient) and non-recurred (n=20, with an average of 65 nuclei per patient) patients. We calculated the features ‘area’, ‘major axis length’, ‘minor axis length’, and ‘roughness’. The p values indicate the t-test results to highlight statistically significant differences between the features of recurred and non-recurred patients.

Figure 4 shows examples of tissue areas from recurrence and non-recurrence patients in which we visualize differences in these morphological features. The elevated values of the nuclei area and roughness parameters suggest that patients who relapsed have bigger nuclei and more nuclei with irregular-shaped cells.

**Figure 4.**
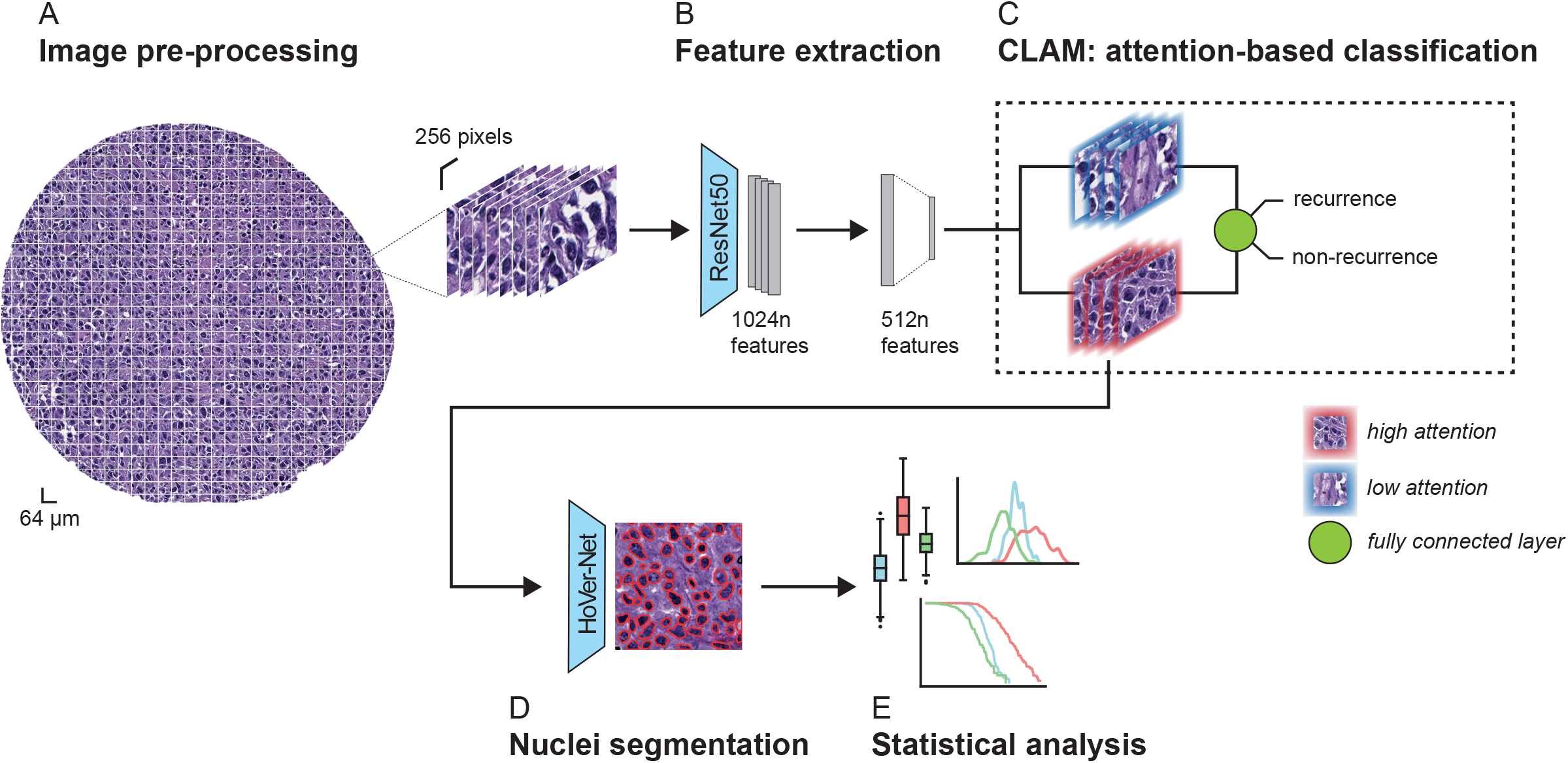
Heatmap visualization of feature values. For the top two patches of recurred and non-recurred patients, we visualize the strengths of the features ‘area’, ‘minor axis length’, and ‘roughness’. The darker the color of a cell the higher its respective feature value.

### 3.3 Survival Prediction

We next inspected whether the visual features identified by our predictive model are predictive of patient survival. To this end, we used the overall survival and the follow-up status from the clinical data of the patients to visualize the survival curves with a Kaplan– Meier plot.

Figure 5 indicates that patients predicted to recur have lower survival probabilities. Additionally, the survival curve of patients predicted as recurring closely matches the curve of patients that actually experienced recurrence based on the clinical follow up data.

**Figure 5.**
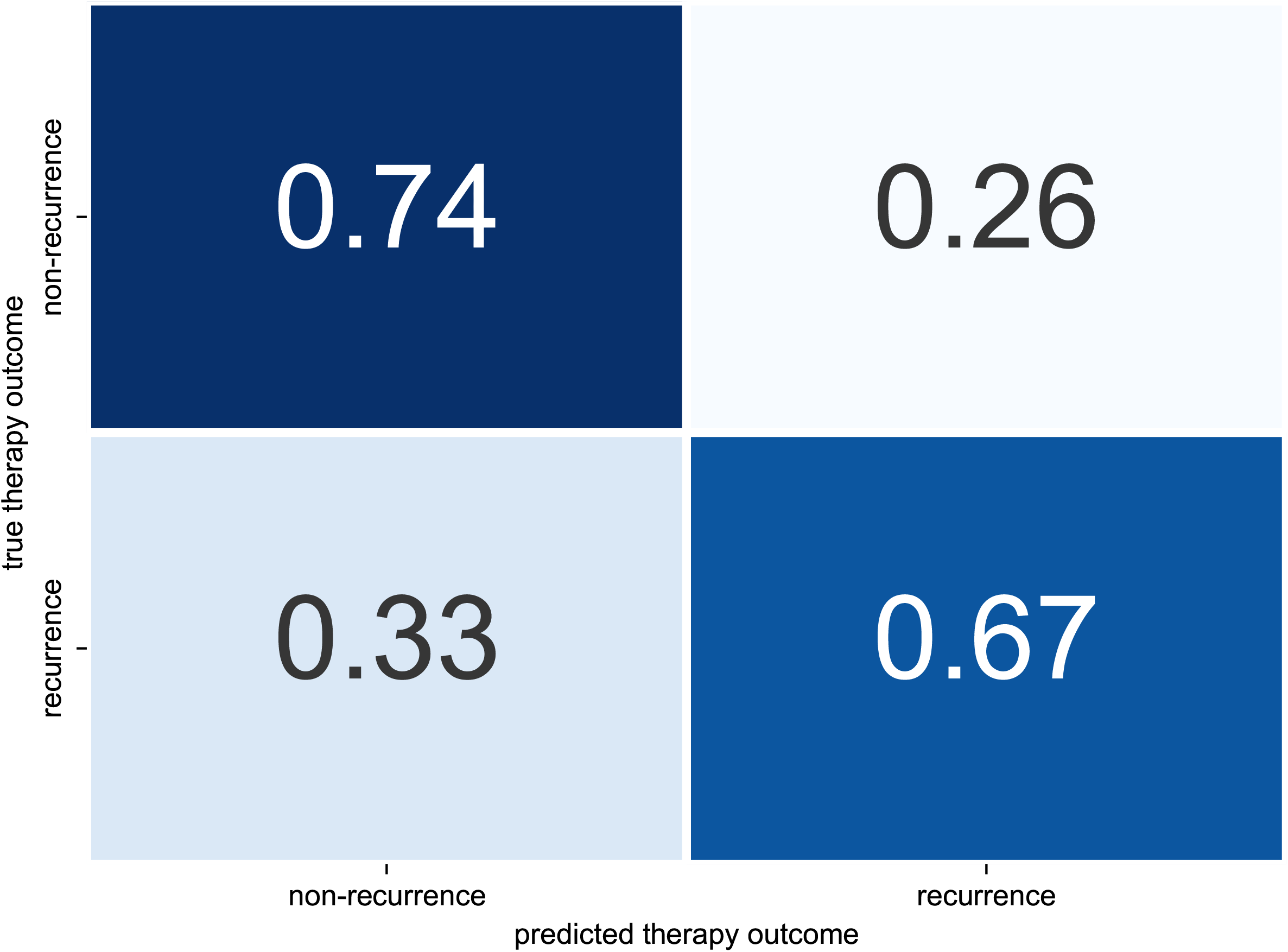
Kaplan–Meier plot for overall survival. The plot shows the overall survival probability of recurred and non-recurred patients (true and predicted) over time. The variable p indicates the t-test-based p-value for true recurrence and non-recurrence and p_pred_ the p value for predicted recurrence and non-recurrence.

## 4 Discussion

Deep learning presents tremendous opportunities for medically relevant prediction tasks, such as recurrence or survival. Additionally, attention-based mechanisms provide insights into the prediction process of the model by enabling quantitative analyses of the tumor morphology and microenvironment features that were crucial to the model’s decision.

In this work, we perform image-based prediction of recurrence of DLBCL. We show, that, despite limited dataset size and class imbalance, our model is able to distinguish recurred from non-recurred patients, indicating that H&E-stained images contain meaningful information for such prediction tasks. Our statistical analysis further suggests that the model’s prediction decision can be explained by morphological characteristics of the cell nuclei within the high-attention areas of the images. Namely, the model focuses on areas where nuclei of recurred patients tend to be bigger and more irregular in shape as the nuclei of non-recurred patients (Figures 3-4). These features indicate a higher degree of nuclear pleomorphism, which has previously been shown as related to tumor aggressiveness [26,27]. Lastly, our model is indirectly predicting patient survival as well (Figure 5).

In addition to the feature extractor pre-trained on ImageNet, we also used the Transformer-based feature extractor CTransPath [28] – a model that has been pretrained on histological images. We found, however, that the model pre-trained on ImageNet yielded better results.

One limitation of our work is the lack of validation on an independent cohort from a different clinical center. To our knowledge, there is however no other publicly available DLBCL histology image data with reported patient clinical outcomes. We aimed to mitigate the model by training and testing the model in several cross validation iterations.

Our study highlights the importance and high information content of histological images for prediction of treatment outcome. In combination with the ability to make our model biologically interpretable, it allows for descriptive, quantitative, and thorough distinction between recurred and non-recurred patients offering the potential for better diagnostics and better understanding of the complexity of fundamental biological processes in aggressive, heterogeneous cancer types, such as DLBCL.

## Authors’ contributions

HN performed data analysis, paper writing and software development. JIP ans SU contributed to software development and data analysis. AS contributed to data analysis. RB and KB designed the study. KB also revised the paper.

## Data availability

All data supporting the results reported in this article can be found here: https://github.com/stanfordmlgroup/DLBCL-Morph.

